# Nitric oxide is important for sensing and survival under hypoxia in Arabidopsis

**DOI:** 10.1101/462218

**Authors:** Aakanksha Wany, Alok Kumar Gupta, Yariv Brotman, Sonika Pandey, Abhay Pratap Vishwakarma, Aprajita Kumari, Pooja Singh, Pradeep K Pathak, Abir U. Igamberdiev, Kapuganti Jagadis Gupta

## Abstract

Nitric oxide (NO) is a free radical molecule that plays an important role in hypoxic stress. We studied the impact of hypoxia-induced NO production on the expression of genes and production of metabolites involved in carbon, oxygen sensing, nitrogen, and antioxidant metabolism using wild type (WT), non-symbiotic haemoglobin-overexpressing (Hb+) and nitrate reductase double mutant (*nia1,2*) of Arabidopsis. Futher application of NO scavenger cPTIO was used to confirm NO role. We found that imposing hypoxia leads to the increaseof NO and reactive oxygen species (ROS) levels in WT, while the reduced levels of NO and higher levels of ROS were observed in roots of Hb*+* and *nia1,2* mutant. Expression of the genes encoding oxygen sensing and the enzymes involved in fermentative pathways, their activities and metabolite levels were highly induced in WT suggesting that NO plays a role in the induction of fermentation. Several genes and metabolites involved in the TCA cycle were also induced in WT in comparison to Hb*+* and *nia* mutant line suggesting that NO can accelerate TCA cycle to regenerate reducing equivalents under hypoxia. Interestingly, we found that the genes and metabolites involved in the ascorbate-glutathione cycle were modulated by NO under hypoxia. The alternative oxidasegene (*AOX1A*) was induced under hypoxia in WTdue to increased levels of NO rather than ROS. Futher, we found that NO improves plant survival. Overall these findings suggest that NO is a major player in plant survival under hypoxia by modulating gene expression and metabolite levels of carbon, nitrogen, oxygen sensing and antioxidant metabolism.

**Highlight:** Hypoxia-induced nitric oxide plays a central role in activation of genes and metabolites involved in fermentation, TCA cycle, nitrogen and antioxidant metabolism

## Introduction

Oxygen is critical for life as it acts as a terminal electron acceptor in mitochondrial electron transport chain, where ATP generation takes place for cellular metabolism, moreover, this process leads to regeneration of NAD^+^ from NADH which can be useful for accelerating fermentation under hypoxia. The importance of oxygen is documented in various cellular pathways such as haem, sterol, amino acid and lipid biosynthesis (Geigenberger, 2003). Unlike animals, plants do not possess sophisticated mechanisms for delivery of oxygen hence, they experience hypoxia during their development due to very low internal oxygen concentrations (Gupta *et al.,* 2009; 2014; Rolletschek *et al.*, 2002). Transient and severe flooding, waterlogging or increased oxygen-consuming microbial activity in soils leads to hypoxia and eventually anoxia (Ricard *et al.,* 1994). During hypoxia, cytochrome c oxidase (COX) activity is ceased that leads to reduced levels of ATP. Under hypoxia, plants produce during oxidation of one hexose only less than one tenth of ATP produced under normoxia but the flux of fermentation can exceed the aerobic metabolic rate and this acceleration leads to production of energy. In addition to energy production, fermentation leads to production of ethanol and lactate (Drew, 1997), and may result in production of harmful levels of reactive oxygen species (ROS) even at low oxygen levels due to the increased redox state (Vergara *et al*., 2012). Severe reduction in root biomass and the allocation of root metabolites to the shoot was also documented (Huang *et al*. 1994).

Plants have several strategies to cope with hypoxic stress. These include various physiological, biochemical and anatomical changes (Bailey-Serres *et al*., 2012). Among the anatomical adaptations, internode elongation (Hattori *et al*., 2009), petiole elongation in Arabidopsis (Mustroph *et al*., 2010; Colmer & Voesenek, 2009), adventitious root formation (Mergemann and Sauter, 2000); lysigenous aerenchyma formation (Abiko *et al*., 2012; Wany *et al.,* 2017), leaf gas films clinging to the surface of leaves of many semi-aquatic species (Winkel *et al*., 2011) were reported. Several metabolic changes were also reported in response to oxygen deprivation (Geigenberger, 2003). Fermentation is combined with some partial activity of the TCA cycle which can help in coping hypoxia (Fox *et al*., 1994). Low oxygen alters gene expression, energy consumption, cellular metabolism and growth. In particular, cells of aerobic organisms have evolved adaptive responses to compensate for the energy loss caused by oxygen deprivation (Gupta *et al.,* 2009, 2014; Rolletschek *et al.*, 2002; Zabalza *et al*., 2009). Examples of such metabolic adaptations to hypoxia include the down-regulation of storage metabolism and the shift from invertase to sucrose synthase pathways of sucrose utilization (Geigenberger, 2003).

Under hypoxic stress, it has been shown that plants produce high amount of nitric oxide (NO), a free radical which plays a role in wide range of processes that include seed germination, stomatal function, root development, cell wall elongation, senescence and programed cell death (Mur *et al*., 2013). In plants, NO can be generated via multiple pathways, those include oxidative and reductive reactions. In the oxidative pathways, NO is produced via oxidation of arginine or hydroxylamine, while the reductive pathways operate via reduction of nitrate and nitrite. The oxidative pathways include NO synthase-like activity (Corpas *et al*., 2009), the polyamine-mediated pathway (Tun *et al*., 2006), and the hydroxylamine-mediated pathway (Rumer *et al*., 2009), these pathways are active under normoxic conditions (Igamberdiev and Hill, 2004; Gupta *et al.,* 2011). NO production from oxidative pathways still needs more characterization. In contrast, the reductive pathways such as the nitrate reductase (NR) pathway (Sakihama *et al*., 2002; Planchet *et al*., 2005), the mitochondrial electron transport pathway (Planchet *et al*., 2005; Gupta *et al.,* 2011), and the plasma membrane nitrite–NO pathway (Stohr *et al*., 2001) operate in response to the decrease in oxygen availability.

Under hypoxic conditions, nitrite reduction to ammonia is inhibited, therefore the accumulated nitrite can act as a substrate for NO production. It was shown that nitrite is a limiting factor for NO production (Planchet *et al*., 2005). Some amount of nitrite is also transported into mitochondria where the complexes III and IV of the electron transport chain produce NO (Gupta *et al*., 2005) which is then scavenged by the cytosolic non-symbiotic haemoglobins (nHb1) via operation of haemoglobin-nitric oxide cycle. So far the function of hypoxically generated NO is attributed to oxidation of NADH (Igamberdiev *et al*., 2006) and generation of limited amount of ATP in roots and nodules (Stoimenova *et al*., 2007; Horchani *et al.,* 2011). NO is also involved in prolongation of plant respiration under hypoxia (Borisjuk *et al*., 2007; Benamar *et al*., 2008) and reduction of ROS via induction of alternative oxidase (AOX) (Gupta *et al.,* 2012; Royo *et al*., 2015).

It was found that NO production initiates within minutes of hypoxia stress (Gupta *et al*., 2005) suggesting that NO could play a role in the induction of genes for hypoxic tolerance. There are many reports suggesting that various genes are induced under hypoxia (Bailey-Serres *et al*., 2012) but, so far no reports are available on how hypoxia-induced NO modulates various genes. Hence, it is very important to check gene expression in the presence of high and low levels of NO therefore, here we investigated the expression of genes induced by NO using WT, Hb+, *nia1,2*. NO donors and NO releasing compounds have potential side effects therefore, we used non-symbiotic haemoglobin overexpressing (Hebelstrup *et al*., 2006) and nitrate reductase double mutant plants that have reduced levels of NO to study the effect of decreased levels of NO on gene induction. Even though NO scavengers such as cPTIO might have side effects but it could be used as complementary to the use of transgenic lines. Therefore, to attribute the observed differences to the effect of NO we have grown WT plants on 100 μM cPTIO. We found that, NO plays an important role in the induction of genes encoding the enzymes of glycolysis, fermentation, TCA cycle and antioxidant metabolism, and that the observed pattern of gene expression changes are correlated with the levels of corresponding metabolites suggesting that NO plays an important role in hypoxia tolerance in plants.

## Materials and methods

### Plant material and growth conditions

The seeds of WT (Col-0) and non-symbiotic hemoglobin (Hb+) over-expressing Arabidopsis (described in Hebelstrup and Jensen, 2008), nitrate reductase double mutant (*nia1,2*); NASC ID N2356, Arginine t-RNA arginyltransferase 1,2 (*ate1,2*); SALK_023492C and proteolysis 6 (*prt6-5*); SALK_051088C mutants were thoroughly rinsed in sterile distilled water for 2-3 times. Surface sterilization was performed by treating the seeds with 1.5 % NaOCl for 7 minutes and thereafter rinsing in sterile autoclaved distilled water for 7-8 times. The seeds were kept in cold chamber for 24 h for stratification and then transferred under sterile conditions on vertical MS agar plates. Plants were grown in a 16 h day: 8 h night regime at 23°C using Phillips TLD 36W 1/830 light bulbs (150 μmolm^-2^ s^-1^ PAR) in Arabidopsis growth room for 21 days. After 21 days, plants were treated with normal air containing 21% of O_2_ (normoxia) or 0.8% oxygen and 99.2% nitrogen gas for 24 h (hypoxia) using Bioxia 800 chamber (IMSET, India). Plants were carefully removed from plates and immediately shock frozen in liquid nitrogen. Results are based on three independent experiments.

### Nitric oxide measurement by DAF fluorescence

For DAF-FM fluorescence, roots were excised from plates immediately after hypoxia treatment (while plates are in hypoxia chamber) and incubated in 10 μM DAF-FM DA (Molecular Probes, Life Technologies, USA) for 15 minutes under dark conditions. To demonstrate that NO was measured, 200 μM of the NO scavenger cPTIO (2-4- carboxyphenyl-4,4,5,5-tetramethylimidazoline-1-oxyl-3-oxide (potassium salt) was used as a control. Then, the root samples were washed three times with the HEPES buffer, transferred to a slide, and mounted with a drop of 10% glycerol with a coverslip. The formation of DAF-2T following NO reaction with DAF-FM DA was visualized using a fluorescence microscope (Nikon*80i*, Japan) with a charge-coupled device camera (DXM1200C) equipped with imaging software (NIS elements). Fluorescence was observed at 495 nm excitation and 515 nm emission. Images were analysed using the Image J software (ver 3.2).

### Measurement of total reactive oxygen species

Total ROS was measured according to the protocol given in Wany *et al.* 2017. Roots were cut with a sharp razor blade into fine segments which were incubated in 10 μM H2-DCFDA (2′,7′-dichlorofluorescein diacetate) for 5-7 minutes at room temperature under dark conditions. The samples were visualized in the fluorescence microscope (Nikon *80i*, Japan; 488 nm excitation and 525 nm emission). Images were analysed using the Image J software (version 3.2).

### Design of primers for carbon, nitrogen and antioxidant pathway genes

The CDS sequences of carbon, nitrogen and antioxidant pathway genes were downloaded from TAIR (The Arabidopsis Information Resource) database and their gene-specific primers were designed using the online available Primer3 tool (ver 0.4.0). The list of primers used to study the expression profiles of above pathway genes are given in the Supplementary Table 1.

### Total RNA isolation, cDNA preparation, and Real-time quantitative reverse-transcription PCR

Total RNA was isolated from roots of 21 days old seedlings using Tri Reagent (MRC, USA) according to the manufacturer’s instructions with slight modifications. Two micrograms of total RNA of each sample was used to synthesize first-strand c-DNA by oligo (dT) 18 primer using Revert Aid™ H-Minus First strand cDNA synthesis kit (Fermentas, Thermo Fisher Scientific, USA). Real-time polymerase chain reaction amplification was carried out using the ABI Prism 7900 sequence detection system (Applied Biosystems), with a power SYBR-green master mix (Applied Biosystems, USA), and the relative quantification of each individual gene expression was performed using the comparative threshold cycle method, as described in the ABI PRISM 7900 Sequence Detection System user bulletin number 2 (Applied Biosystems, USA). Actin, 18srRNA and tubulin was used as internal references. For data normalization, 18sRNA was considered. The amplification program used for quantitative reverse-transcription polymerase chain reaction was 95 °C for 2 min, 40 cycles at 95 °C for 30 s, 60 °C for 30 s, 72 °C for 30 s, 60 °C for 15 s, and 95 °C for 15 s. All samples were amplified in triplicates, and the mean Ct values were considered.

### GC-MS analysis

GC-MS analysis was performed as described previously (Lisec *et al.*, 2006). Five biological replicates were used for analysis. A target search software package was used to identify the peaks (Cuadros-Inostroza *et al.*, 2009). Metabolites were selected by comparing their retention indexes (± 2 s) and spectra (similarity >85%) against the compounds stored in the Golm Metabolome Database (Kopka *et al.*, 2005). Metabolite data were log10-transformed to improve normality as described by Steinfath *et al.* (2008) and the data were normalized to show identical medium peak sizes per sample group. Day-normalization and sample median-normalization were conducted; the resulting data matrix was used for further analysis (Wu *et al.,* 2016).

### Nitrate reductase assay

Seedlings (1g) were harvested and immediately finely ground with liquid nitrogen. The tissue powder was homogenized in 2 ml ice cold extraction buffer containing 100 mM HEPES pH 7.6, 3.5 mM mercaptoethanol, 15 mM MgCl2, 0.5% PVP, 0.5% BSA and 0.3% of Triton X100.The homogenate was centrifuged at 14000 rpm at 4°C for 30 minutes to obtain a clear supernatant. This supernatant was used to determine NR assay. Prior to NR activity measurement, the protein levels were measured by Bradford’s method. 100 μg of total protein were supplied with the NR assay buffer containing 50 mM HEPES-KOH pH 7.6, 5 mM KNO3 and 0.2 mM NADH in a total volume of 800 μl. After incubating this reaction mixture for 15 min (RT), the reaction was stopped by adding 125 μl of zinc acetate (0.5 M) and Griess reagent [0.5 ml sulfanilamide and 0.5 ml N-(1-naphthyl) ethylenediamine dihydrochloride solution] available in the Griess reagent kit (Molecular probes, Inc., Invitrogen, Life Technologies, USA). The nitrite content was determined colorimetrically at 546 nm. The control reaction was carried out using 10 μM potassium nitrite solution in order to check colour development with the Griess reagent (Planchet *et al.*, 2005).

### ADH assay

Roots of 21 days old seedlings (0.5 g) were harvested and finely ground in liquid nitrogen. The tissue powder was homogenized in 500 μl of ice-cold extraction buffer containing 50 mM HEPES pH 7.5, 15% (v/v) glycerol, 1 mM EDTA, 3 mM MgCl2, 10 mM dithiothreitol (DTT) and 2 mM phenylmethylsulphonyl fluoride (PMSF). It was repeatedly vortexed for 3 times until complete solubilization of the tissue. The samples were then centrifuged at 14000 rpm, 4°C for 20 min and the clear supernatant was transferred into a new tube. This crude protein extract was used to determine ADH activity. First, the protein levels were measured by Bradford’s method (Bradford, 1976). 100 μg of total protein were supplied with the assay buffer containing 50 mM sodium phosphate buffer pH 8.0, 300 μM NAD^+^ and 150 mM ethanol in a total volume of 200 μl. Then, the specific ADH activity was measured in a 96-well microplate (96-well U-Bottom, clear PS microplate, Greiner-Bio-One, Sigma-Aldrich, India) set in the kinetic mode at discrete wavelength of 340 nm in the microplate reader (Polar Star Omega, BMG LabTech, Germany). The reaction was followed by measuring the increase in absorbance at 340 nm. The specific ADH activity was calculated as units per μg protein, i.e., μmol amount of NADH formed per minute and per μg total protein, with an extinction coefficient of ε = 6200 l mol^-1^ cm^-1^.

### Protein extraction and immunoblot analysis

Frozen Arabidopsis seedlings (0.5 g) were crushed to fine powder in liquid nitrogen and mixed well in 100 μl ice-cold extraction buffer containing 50 mM Tris HCl buffer (pH 7.2), 3.5 mM 2-mercaptoethanol, 1 mM PMSF and 15 mM MgCl2. Homogenized samples were vortexed vigorously three times at one minute interval each and centrifuged at 14,000 rpm for 15 min at 4°C. Supernatant contained the extracted protein, was quantified by Bradford’s assay (Bradford, 1976) using 1 mg/ml BSA as standard. Protein samples (50 μg) were mixed with sample loading buffer and after denaturation, were fractionated on a 12% polyacrylamide gel. Protein from gel was transferred to nitrocellulose blotting membrane (Pall Corporation, Florida) in a wet transblot system (BIORAD, India) according to manufacturer’s protocol for 2.30 h at 50V. Membrane was blocked with 5% (w/v) skimmed milk in Tris-buffered saline (TBS) buffer for 1 h, and kept for incubation on rotary shaker in primary antibody; polyclonal anti-AOX1,rabbit serum,(1:1000 dilution, Agrisera) at 4°C. After that, membrane was washed in TBS-T (TBS-Tween 20) for three times and was incubated with secondary antibody, polyclonal goat anti-Rabbit IgG (H + L) horseradish peroxidase conjugate, (1:5000 dilution, Pierce Protein Biology) in TBS-T at room temperature for 1.30 h. Again, membrane was washed for three times after every 10 min interval with TBS-T. For protein detection, membrane was incubated for 5 minutes in dark in ECL mix (Clarity™ Western ECL Blotting Substrates, BIORAD, India) in a ratio of 1:1 of solution A and B according to the manufacturer’s instructions. The membrane was then visualized in ChemiDoc MP Imaging system (10022684, BIORAD, India).

## Results

### Hypoxic stress leads to high increase in NO levels and moderate increase in ROS levels

First, we tested the impact of hypoxic stress on induction of NO and ROS levels in WT and Hb+ and *nia1,2* plants of Arabidopsis. For this purpose, we have grown plants on Murashige and Skoog (MS) medium agar plates in vertical orientation for three weeks and then we placed these plates in a hypoxia chamber set at 0.8% oxygen and flushed for 24 h. Immediately after hypoxia treatment, we checked for NO levels using DAF-FM DA and found that under hypoxia, WT plants produced significant amount of NO in comparison to Hb+ and *nia1,2* mutant (Fig 1A&B). NO scavenger cPTIO effectively scavenged DAF-FM DA fluorescence (Fig 1A right panel & Fig1B). In parallel, we measured total ROS levels by incubating roots in H2DCF DA and measured DCF fluorescence. As shown in Fig 1C, Hb+, *nia1,2* mutant roots produced slightly higher levels of ROS under normoxia in comparison to WT; but under hypoxia, ROS levels have increased in all genotypes, while higher levels of ROS were observed in Hb+ and *nia1,2* mutant in comparison to WT roots suggesting that hypoxia-induced NO can control ROS levels.

**Fig 1:**
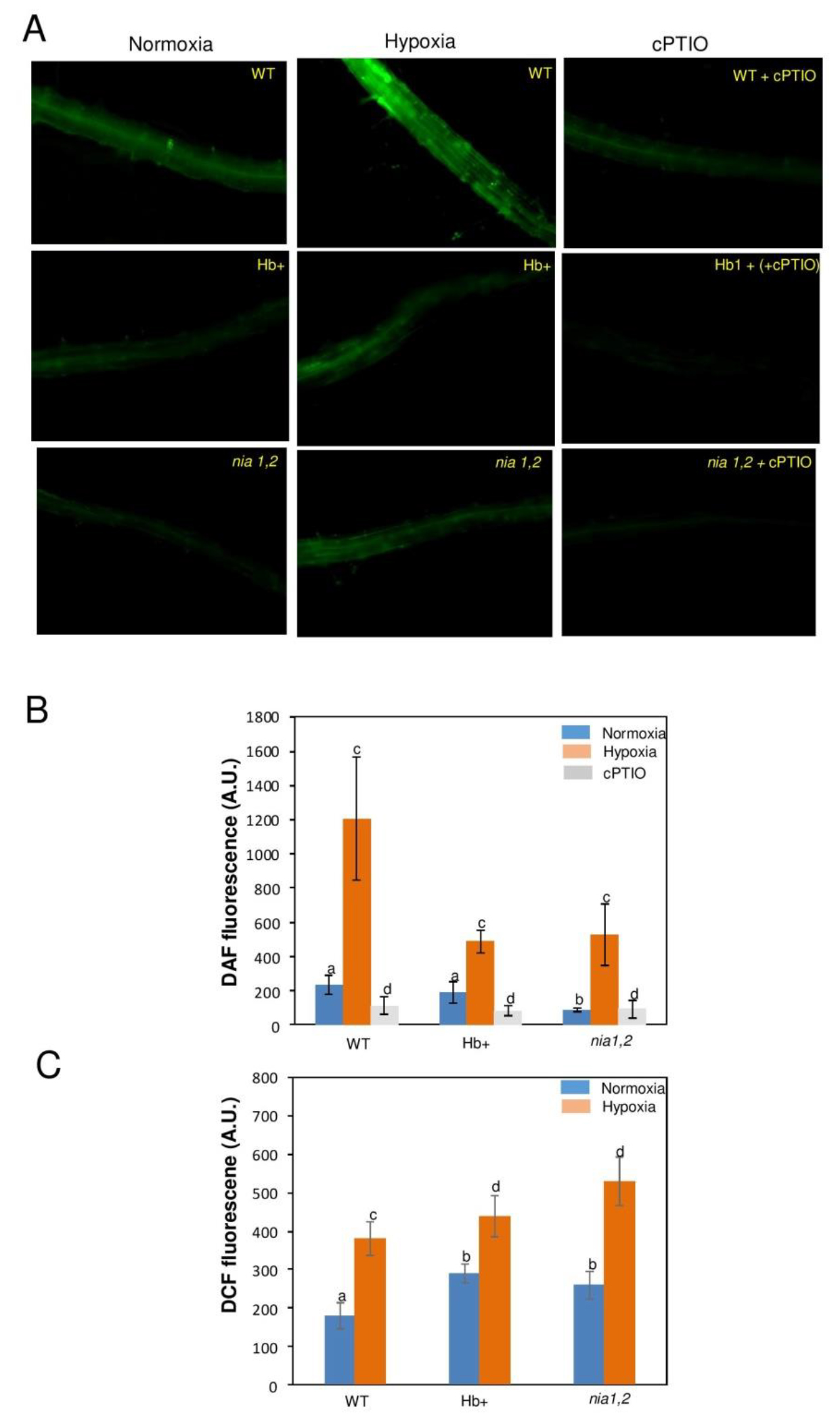
**A**: DAF FM fluorescence from WT, Hb+ and *nia 1,2* mutant roots under normoxia and hypoxia 24 h, treated with 10 μM DAF-FM DA and 200 μM cPTIO + 10 μM DAF FM DA. Images are representative of 3 independent replicates. **B**: NO production by WT and Hb+ and *nia1,2* roots under normoxia and hypoxia quantified by Image J. Fluorescence from hypoxia-treated roots with 200 μM cPTIO + 10 μM DAF-FM DA used as a control. Graphs show one representative measurement of 8 biological replicates. Error bars indicate SD. Means (n=4) with different letters are significantly different (one-way ANOVA, p< 0.05) **C**: Reactive oxygen species (ROS) levels visualized by 2′,7′-dichlorofluorescein (H_2_DCFDA) from *Arabidopsis* roots of WT, Hb+ and *nia1,2* mutant under normoxia and hypoxia 24 h. Means (n=4) with different letters are significantly different (one-way ANOVA, p< 0.01)

### Genes involved in nitrogen metabolism are differentially modulated by nitric oxide under hypoxia

NO measurements under hypoxia revealed that WT roots produce higher levels of NO in comparison to Hb+ & *nia1,2* mutant roots (Fig 1A&B). Then, we monitored the impact of higher levels of NO on the genes involved in nitrogen metabolism (Fig 2). First, we tested the expression of *NITRATE REDUCTASE* gene (*NIA1)* which is responsible for NO production (Bright *et al.,* 2006). We found 3-fold increase in *NIA1* expression in response to hypoxia in WT whereas in all other genotypes it was not at all induced (Fig 2A) The *NIA1* expression has correlated with NR activity where 2.5 fold elevation was observed in WT under hypoxia (Fig 2F). This can indicate that an increased level of NO is mainly due to nitrate reductase induction and increased NR activity. NR catalyses nitrate to nitrite conversion and the produced nitrite is reduced to ammonium in plastids which is then assimilated to amino acids via GS/GOGAT cycle. Therefore, we tested the expression of cytosolic *GLUTAMINE SYNTHETASE-1*(*GLN1*). We found increased levels of *GLN1* expression in WT under normoxia, whereas the levels were suppressed in Hb+, *nia* mutant *and* cPTIO-grown plants under normoxia (Fig 2B). Under hypoxia, 2- fold increase was observed in WT, whereas in Hb+, the expression has decreased 2- fold in comparison to WT under hypoxia. This gene was almost not induced in *nia1,2* mutant and in cPTIO grown plants. The levels of the main metabolic product of GS/GOGAT pathway, glutamine, were also elevated under hypoxia (Fig 8a). Expression profile of *FERREDOXIN-DEPENDENT GLUTAMATE SYNTHASE* (*GLU2; fd-GOGAT2*) gene revealed that under normoxia, WT showed its higher expression than Hb+, *nia* mutant and cPTIO-grown plants while in the plants imposed to hypoxia, 2- fold induction was observed in WT, whereas in Hb+, *nia1,2* and cPTIO-grown plants it was downregulated in comparison to WT under hypoxia (Fig 2C). 2-oxoglutarate (2-OG) levels were decreased in WT under hypoxia while the increased levels were observed in Hb+ (Fig 8b).Glutamate levels were also higher in WT under hypoxia (Fig 8o). Then, we checked expression levels of *ALANINE AMINOTRANSFERASE 1* gene (*AlaAT1)* which catalyses the reversible transfer of an amino group from glutamate to pyruvate to form 2-OG and alanine. We found approximately, 2 fold induction of this gene in WT under hypoxia, whereas in Hb+, *nia1,2* and cPTIO grown plants it was downregulated in comparison to WT under hypoxia (Fig 2D). Increased levels of alanine were also observed under hypoxia in WT (Fig 8c) and then, another important gene that is involved in amino acid biosynthesis is *2-OXOGLUTARATE DEHYDROGENASE* (*OGDH)* which links carbon and nitrogen metabolism. Expression profile indicated that *OGDH* expression has decreased in both WT, Hb+, *nia1,2* and cPTIO grown plants under hypoxia (Fig 2E).

**Fig 2:**
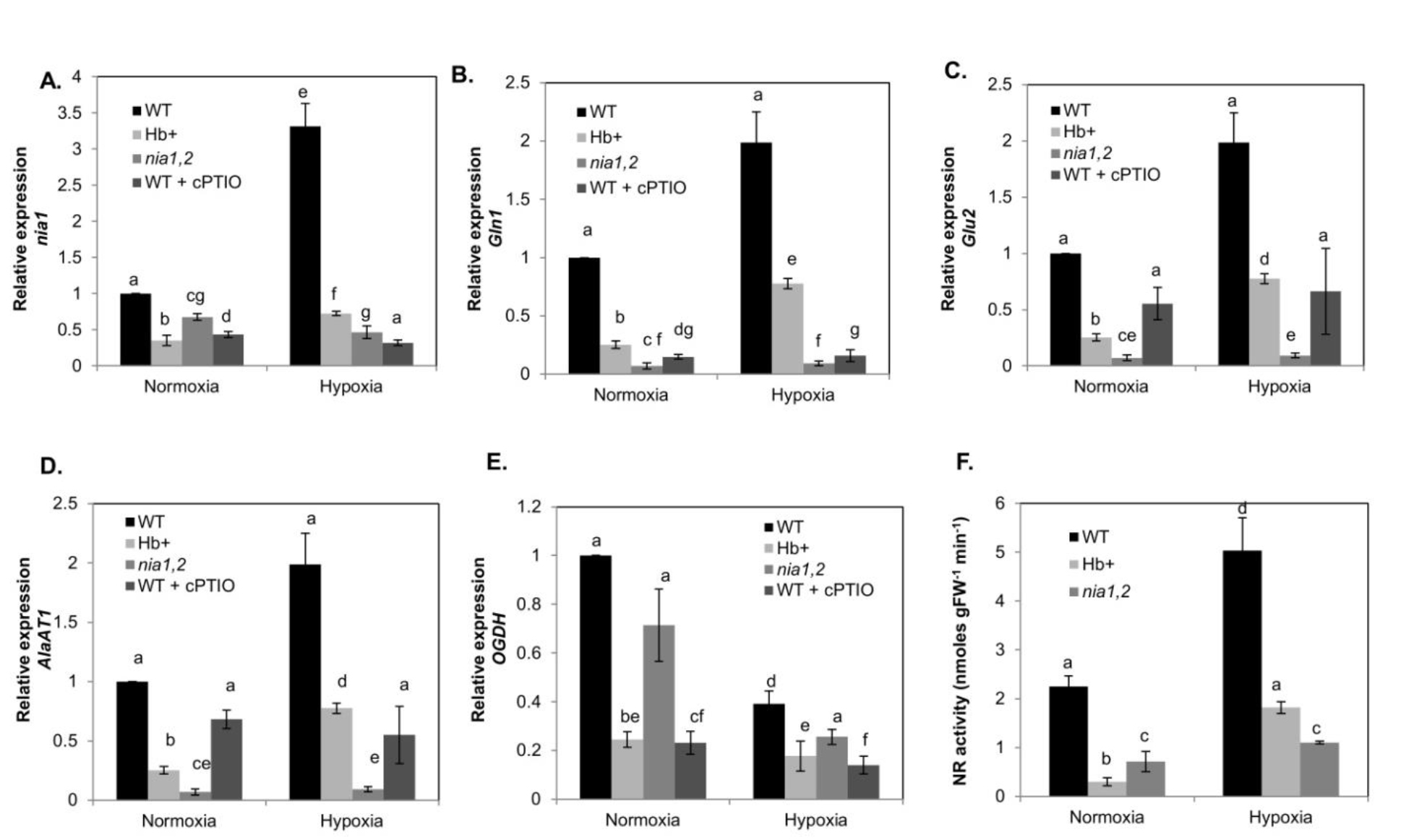
Relative expression of genes involved in nitrogen metabolism: (A) *NIA1* (*NITRATE REDUCTASE1*; AT1G77760), (B) *Gln1* (*GLUTAMINE SYNTHETASE-1;* AT1G48470), (C) *GLU2 (GLUTAMATE SYNTHASE-2*; AT2G41220), (D) *ALaAT1 (ALANINE AMINOTRANSFERASE-1*; AT1G17290) and (E) *OGDH-E1 (2- OXOGLUTARATE DEHYDROGENASE, E1 COMPONENT*; AT3G55410) in wild-type (WT-black bars), non-symbiotic hemoglobin-1 over-expression lines (Hb+– light grey bars), nia double mutants (*nia1,2*– grey bars) and cPTIO pre-treated WT plants (dark grey bars) under normoxia and hypoxia-24h (0.8% O2) (F). NR activity from total seedlings of WT, Hb+, and *nia1,2* mutants under normoxia and hypoxia-24h. Values are means (n=3) ± SE. Statistical significance between normoxia and hypoxia-24h treatments is evaluated by the analysis of variance (ANOVA) followed by different letters indicating significant difference according to Tukey HSD at p<0.01.

### Genes involved in glycolysis and fermentation are highly expressed in response to hypoxia-induced NO

Glycolysis and fermentative metabolism play a role in survival of plants under hypoxia via recycling of NAD^+^, therefore, it is essential to know whether NO modulates these pathways. We checked the expression of sucrose synthase which is a key enzyme involved in catabolism of sucrose for mobilising it for plant growth, development and stress. Since operation of glycolysis requires glucose, the process depends on *SUCROSE SYNTHASE 1* (*Sus1*) expression. Our results show that *Sus1* gene displayed very low levels of expression under normoxia in Hb+, *nia* mutant and cPTIO-grown plants. Four times elevated expression was observed in WT under hypoxia, whereas in Hb+, *nia* mutant and cPTIO-grown plants the expression levels were 3-4 times less than in WT under hypoxia (Fig 3A) suggesting that NO modulates *Sus1* expression under hypoxia which is essential for accelerating glycolysis. Sucrose levels also increased under hypoxia in WT (Fig 8e). Pyruvate decarboxylase (PDC) converts pyruvate to acetaldehyde which can be used by alcohol dehydrogenase (ADH) to produce ethanol that leads to regeneration of NAD^+^. Therefore, we checked expression of *PDC1* and *ADH1* (Fig 3B and C). Under normoxic condition, we observed down-regulation of *PDC1* in Hb+ and *nia* roots, whereas under hypoxia, its expression has increased up to 2.2 fold in WT whereas moderate induction (but less than WT) was observed in Hb+, *nia* and cPTIO grown plants. *ADH1* gene expression was also upregulated nearly 5-fold under hypoxia in WT, 3.6-fold in Hb+ lines and 2- fold in *nia1,2* mutant. A similar pattern of increase was observed in ADH activity where a four times increased activity was observed in WT under hypoxia, 2-fold increase in Hb+ and 3-fold increase in *nia1,2* mutant (Fig 3F) *LACTATE DEHYDROGENASE* (LDH) also recycles NAD^+^ via conversion of pyruvate to lactate. Under hypoxia, *LDH* gene expression has increased upto 1.3-fold in WT, whereas no induction was observed in both Hb+ line, *nia1,2* mutant and cPTIO-grown plants (Fig 3D). Pyruvate kinase catalyzes the final step of glycolysis by the transfer of a phosphate group from phosphoenolpyruvate (PEP) to adenosine diphosphate (ADP), yielding one molecule of pyruvate and one molecule of ATP. *PYRUVATE KINASE* gene expression increased up to 3.5-fold in WT roots whereas no induction was observed in Hb+ lines, *nia1,2* mutant and cPTIO-grown plants upon imposing hypoxia (Fig 3E). PEP levels have slightly increased under hypoxia in WT, significantly increased Hb+ but decreased in *nia1,2* mutant. Pyruvate levels have increased higher in WT under hypoxia whereas in Hb+ and *nia1,2* mutant no change was observed (Fig 8d)

**Fig 3:**
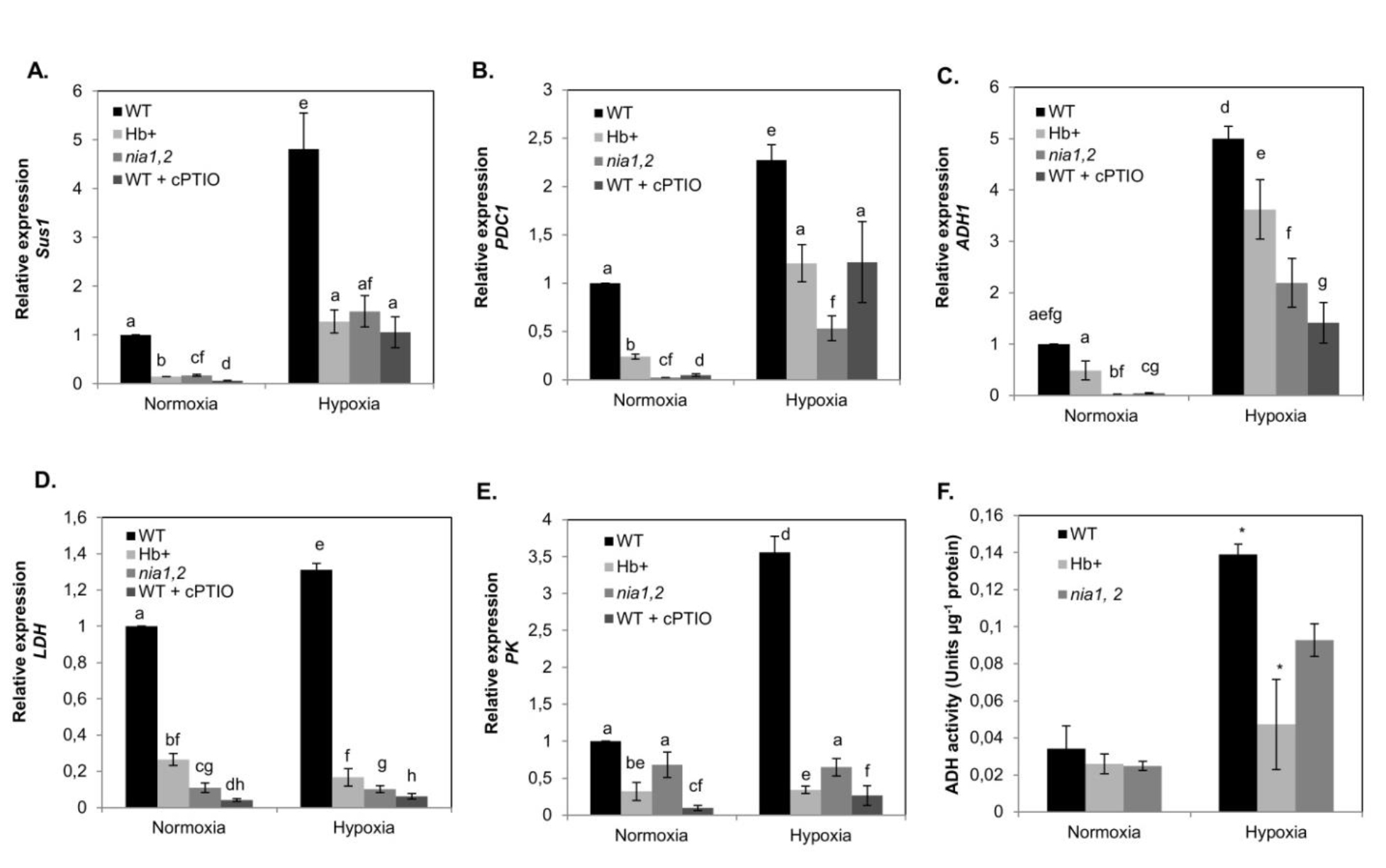
Relative expression of genes involved in carbohydrate metabolism and fermentative pathways: (A) *Sus1* (*SUCROSE SYNTHASE-1*; AT5G20830), (B) *PK* (*PYRUVATE KINASE;* AT2G36580), (C) *LDH (LACTATE DEHYDROGENASE*; AT4G17260), (D) *PDC1 (PYRUVATE DECARBOXYLASE-1*; AT4G33070) and (E) *ADH1 (ALCOHOL DEHYDROGENASE-1*; AT1G77120) in wild-type (WT– black bars), non-symbiotic hemoglobin-1 over-expression lines (Hb+- light grey bars), nia double mutants (*nia1,2*– grey bars) and cPTIO pre-treated WT plants (dark grey bars) under normoxia and hypoxia-24h (0.8% O2) (F). ADH activity from total seedlings of WT, Hb+, and *nia1,2* mutants under under normoxia and hypoxia-24h. Values are means (n=3) ± SE. Statistical significance between normoxia and hypoxia-24h treatments is evaluated by the analysis of variance (ANOVA) followed by different letters indicating significant difference according to Tukey HSD at p<0.01.

### Genes involved in oxygen sensing are differentially regulated by nitric oxide

Recently it was demonstrated that group VII ERF transcription factors which are involved in hypoxia survival are responsible for induction of genes involved in fermentative metabolism (Gibbs *et al.,* 2011, Licausi *et al.,* 2011). In order to correlate the increased expression of fermentative genes and ADH activity (Fig 3 B,C,D), we checked the expression of these transcription factors and other genes involved in oxygen sensing in WT, Hb+, *nia1,2*, *ate1-2*, *prt6-5* backgrounds (Fig 4).

**Fig 4:**
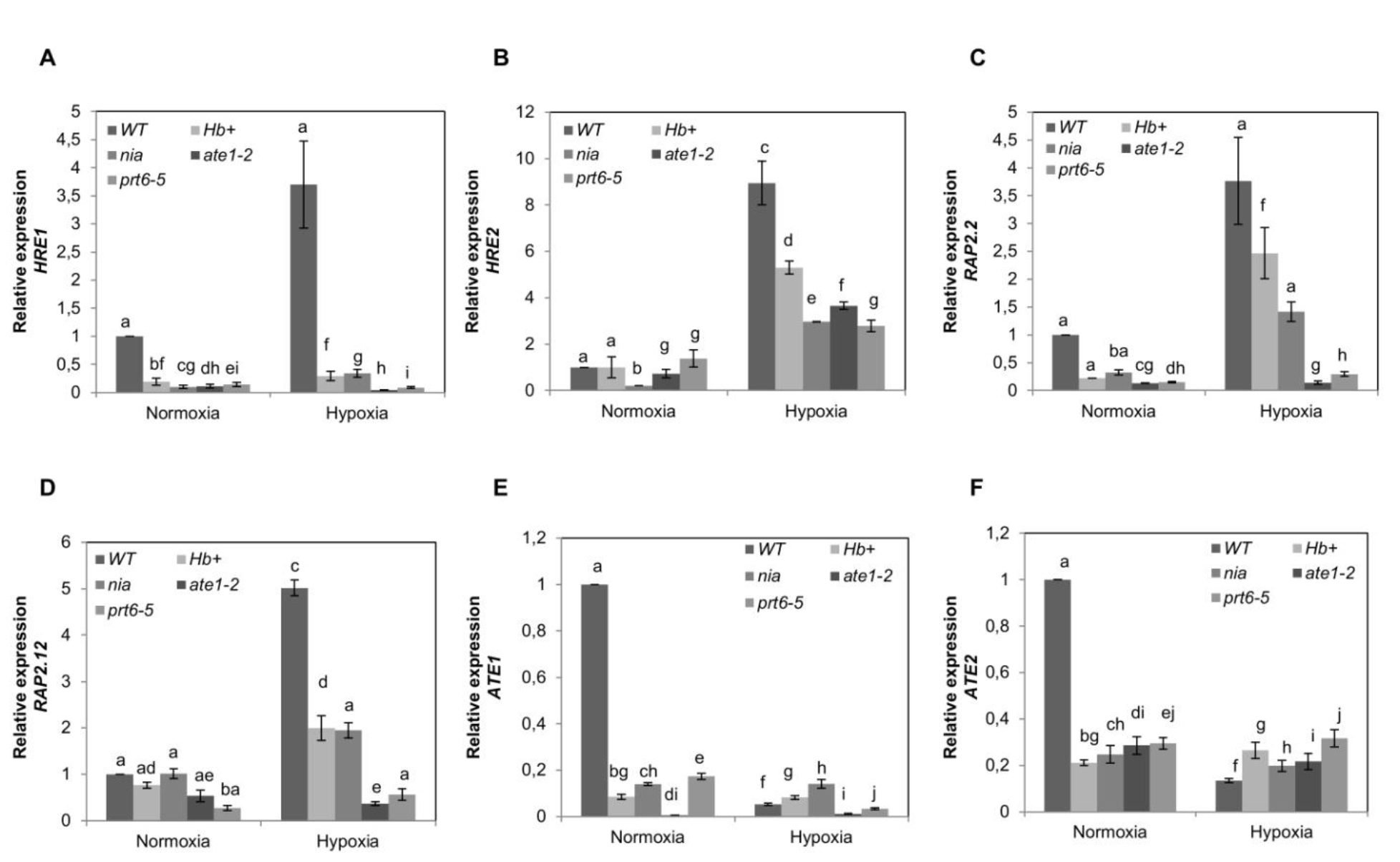
Relative expression of genes involved in oxygen sensing. (a) *HRE1 (HYPOXIA RESPONSE ERF-1*; AT1G72360), (b) *HRE2 (HYPOXIA RESPONSE ERF-2*; AT2G47520), (c) *RAP2.2 (ERF/AP2 TRANSCRIPTION FACTOR FAMILY*; AT3G14230), (d) *RAP2.12 (ERF/AP2 TRANSCRIPTION FACTOR FAMILY*; AT1G53910), (e) *ATE1 (ARGININE-T-RNA PROTEIN TRANSFERASE 1*; AT5G05700), (f) *ATE2 (ARGININE-T-RNA PROTEIN TRANSFERASE 2*; AT3G11240) in wild-type (WT), non-symbiotic hemoglobin-1 over-expression lines (Hb+), *nia* double mutants (nia1,2), *ate1-2 and prt6-5* mutants under normoxia and hypoxia-24h (0.8% O2). Values are means (n=3) ± SE. Statistical significance between normoxia and hypoxia-24 h is analyzed by analysis of variance (ANOVA) followed by different letters indicating significant difference according to Tukey HSD at p<0.01

The expression of gene encoding *HYPOXIA RESPONSE ERF-1 (HRE1)* transcription factor has upregulated 3-4 fold in WT under hypoxia where as in Hb+, *nia1,2* it was not at all induced. We further checked expression of this gene in N-end rule pathway mutants *ate1-2* and *prt6-5*. As expected this gene was not induced in these mutants background (Fig 4A). Another important gene which plays an important role in oxygen sensing *HRE2* was upregulated under hypoxia in WT, Hb+, *nia1,2*, *ate1-2*, *prt6-5* background but in WT, further increased expression was observed (Fig 4B). The expression of *RAP2.2* and *RAP2.12* has increased under hypoxia in WT, Hb+, *nia1,2* but not in *ate1-2* and *prt6-5* mutants (Fig 4C&D). *ARGININE-tRNA PROTEIN TRANSFERASE 1* (*ATE1)* and *ARGININE-tRNA PROTEIN TRANSFERASE 2* (*ATE2*) which have been shown to be involved in oxygen sensing (Licausi *et al.,* 2011) were also checked. Expression of *ATE1* was upregulated in WT under normoxia but this gene was not at all induced in other genotypes under normoxia. Interestingly, this was gene was neither induced in WT nor in other genotypes under hypoxia (Fig 4E). Under normoxia, *ATE2* displayed reduced expression in Hb+, *nia1,2 ate1-2* and *prt6-5* in comparison to WT whereas under hypoxia, also this gene displayed reduced expression with an exception that WT showed slightly lower expression than other genotypes.

### TCA cycle and alternative mitochondrial pathways are affected by hypoxic NO

We further checked expression of genes encoding the enzymes of TCA cycle (Fig 5). First, we checked the expression of *PYRUVATE DEHYDROGENASE* (*PDH*) *gene* encoding pyruvate dehydrogenase which converts pyruvate to acetyl-CoA. We found that the expression levels of *PDH* are significantly down-regulated under hypoxia in WT, Hb+ lines and *nia1,2* mutant (Fig 5A).Then, we observed *CITRATE SYNTHASE* (*CS4*) gene which is an entry enzyme of the tricarboxylic acid (TCA) cycle. This enzyme catalyzes the reaction between acetyl-coenzyme A (acetyl CoA) and oxaloacetic acid to form citric acid. We found almost four fold increase in *CS4* expression in WT, whereas no induction was seen in Hb+, *nia1,2* mutant and cPTIO-grown plants under hypoxia. (Fig 5B). Then, we monitored the expression of *ACONITASE* (*ACO2*) gene which mediates conversion of citrate to isocitrate. Under hypoxia, *ACO2* has more than double expression in WT roots in comparison to its expression under normoxia while there was absolutely no change in *ACO2* expression levels in both Hb+, *nia1,2* mutant and cPTIO grown plants under hypoxia (Fig 5C).The analysis of *SUCCINYL COA SYNTHETASE (SCS),* which encodes the enzyme that converts succinyl-CoA to succinate, revealed that *SCS* expression is severely down-regulated under hypoxia in both WT, Hb+ lines and *nia1,2* mutant (Fig 5D). Increased levels of succinate levels were found in WT under hypoxia but no change was observed in Hb+ and *nia1,2* mutant (Fig 8j).Then, we monitored expression of gene *SUCCINATE DEHYDROGENASE* (*SDH2*) encoding the next TCA cycle enzyme which mediates conversion of succinate to fumarate, this process leads to regeneration of FAD. Under normoxia, the expression of *SDH2* was slightly higher in Hb+ line and *nia1,2* mutant as compared to WT whereas very low expression was observed in cPTIO grown plants. But, upon imposing hypoxia, there was a 5-fold induction of *SDH2* in WT. This gene has significantly downregulated in Hb+ line, *nia1,2* mutant and cPTIO grown plants under hypoxia. Then, we checked the expression of *FUMARASE* (*Fum1*) encoding fumarate hydratase that catalyzes the reversible hydration/dehydration of fumarate to L-malate. We found that, under normoxia its expression is around 2-fold higher in Hb+ line and 2.7 fold higher in *nia1,2* mutant in comparison to WT, but under hypoxia, a ~7 fold increased expression was seen in WT. In Hb+ line, *nia1,2* and cPTIO grown plants this gene was significantly down-regulated. Fumarate and malate levels have increased in WT under hypoxia (Fig 8k&l)

**Fig 5:**
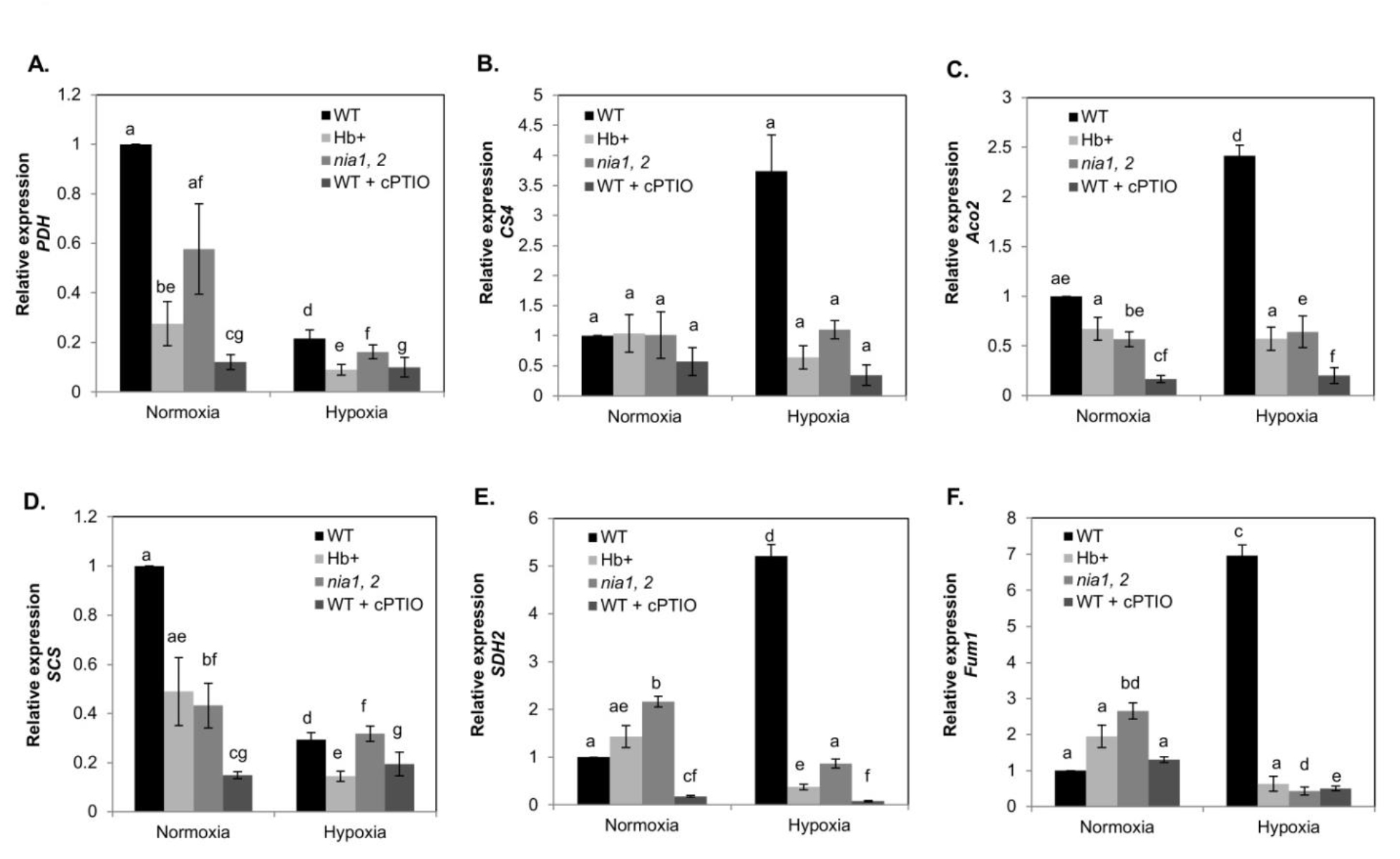
Relative expression of genes involved in TCA pathway: (A) *PDH1 E1 subunit* (*PYRUVATE DEHYDROGENASE-1*; AT1G01090), (B) *CS4* (*CITRATE SYNTHASE-4;* AT2G44350), (C) *ACO2 (ACONITASE-2*; AT4G26970), (D) *SCS* or ACL *(SUCCINYL-COA SYNTHETASE, BETA SUBUNIT OR ATP CITRATE LYASE*; AT2G20420), (E) *SDH2 (SUCCINATE DEHYDROGENASE1-2*; AT2G18450); (F) *FUM1* (*FUMARASE-1*, AT2G47510) in wild-type (WT-black bars), non-symbiotic hemoglobin-1 over-expression lines (Hb+- light grey bars), *nia* double mutants (nia1,2– grey bars) and cPTIO pre-treated WT plants (dark grey bars) under normoxia and hypoxia-24h (0.8% O_2_). Values are means (n=3) ± SE. Statistical significance between normoxia and hypoxia-24h treatments is evaluated by the analysis of variance (ANOVA) followed by different letters indicating significant difference according to Tukey HSD at p<0.01.

### Antioxidant genes are upregulated by hypoxia-induced NO

Antioxidants are very important for protection from the excessive formation of ROS. In order to check whether NO plays a role in inducing antioxidants, we analysed the expression of various antioxidant-related genes (Fig 6&7). First, we checked the expression of *ASCORBATE PEROXIDASE* (*APX1*) gene. This gene encodes the enzyme that detoxifies hydrogen peroxide using ascorbate as a substrate. Expression of *APX1* has increased in Hb+ line nearly ~2 fold under normoxia in comparison to WT. Under hypoxia, there was a marked increase in *APX1* expression up to ~3 fold in WT where as in Hb+ line, *nia1,2 and* cPTIO grown plants it displayed down-regulation (Fig 6A). Then, we checked the expression of *ASCORBATE OXIDASE* encoding another important antioxidant enzyme ascorbate oxidase; which uses ascorbate as a substrate and converts it into dehydroascorbate. Under normoxia, the *ascorbate oxidase* gene expression was 3 fold higher in *nia1,2* mutant in comparison to Hb+ and WT, whereas under hypoxia~5 times induction was observed in WT while in Hb+ line and *nia1,2* mutant, it was strongly downregulated (Fig 6B)*. MONODEHYDROASCORBATE REDUCTASE* (*MDAR1*) encodes the enzyme that recycles oxidised forms of ascorbate, this plays a role in the ascorbate-glutathione cycle. In WT, Hb+ and *nia1,2* mutant, low levels of expression were seen under normoxia whereas under hypoxia, 4 fold increase in WT was observed (Fig 6C). Important metabolites such as ascorbate and dehydroascorbate exhibited increased levels under hypoxia in WT (Fig 8m&n). *GLUTATHIONE REDUCTASE* (*GR1*) is another antioxidant gene in the ascorbate-glutathione cycle. GR enzyme catalyses the reduction of glutathione disulfide (GSSG) to the sulfhydryl form of glutathione (GSH), this contribute to the increased levels of reduced glutathione. Approximately, ~3 fold increase was observed in its expression in WT whereas no change was detected in Hb+ plants under hypoxia. The *GR1* expression levels are induced up to 1.5 fold in *nia1,2* plants under normoxia and slightly more induced under hypoxia (Fig 6D). Expression profile of another antioxidant gene *MANGANESE SUPEROXIDE DISMUTASE 1 (MSD1)* revealed that there was ~2-fold induction in its expression under hypoxia in WT, whereas it was down-regulated in Hb+ lines and *nia1,2* under hypoxia as compared to WT normoxia (Fig 6E). We checked expression of *ALTERNATIVE OXIDASE1A* (*AOX1A*) which encodes an important mitochondrial bypass enzyme and plays a role in stress tolerance (Fig 7A).There was a trend towards decrease of *AOX1A* expression in Hb+ lines and *nia1,2* under normoxia, but under hypoxia, there was ~4 fold increase in its expression in WT and very slight induction in Hb+ lines and 1.5 fold in *nia1,2* under hypoxia as compared to normoxia (Fig 7A).The increased expression of *AOX1A* could be due to increased citrate levels (Fig 7h). It was previously shown that increased citrate levels directly induce AOX capacity and protein levels under hypoxia (Gupta *et al.,* 2012). Increased AOX1A protein levels were observed WT and reduced levels were observed in Hb+ and *nia1,2* mutant under hypoxia (Fig 7C) The mitochondrial externally facing *NADPH DEHYDROGENASE 1* (*NDB1*) expression was slightly increased in Hb+ lines under normoxia as compared to WT. But, upon imposing hypoxia, ~5 fold induction was observed in WT, and absolutely no change was detected in Hb+, *nia1,2* mutant and cPTIO grown plants.

**Fig 6:**
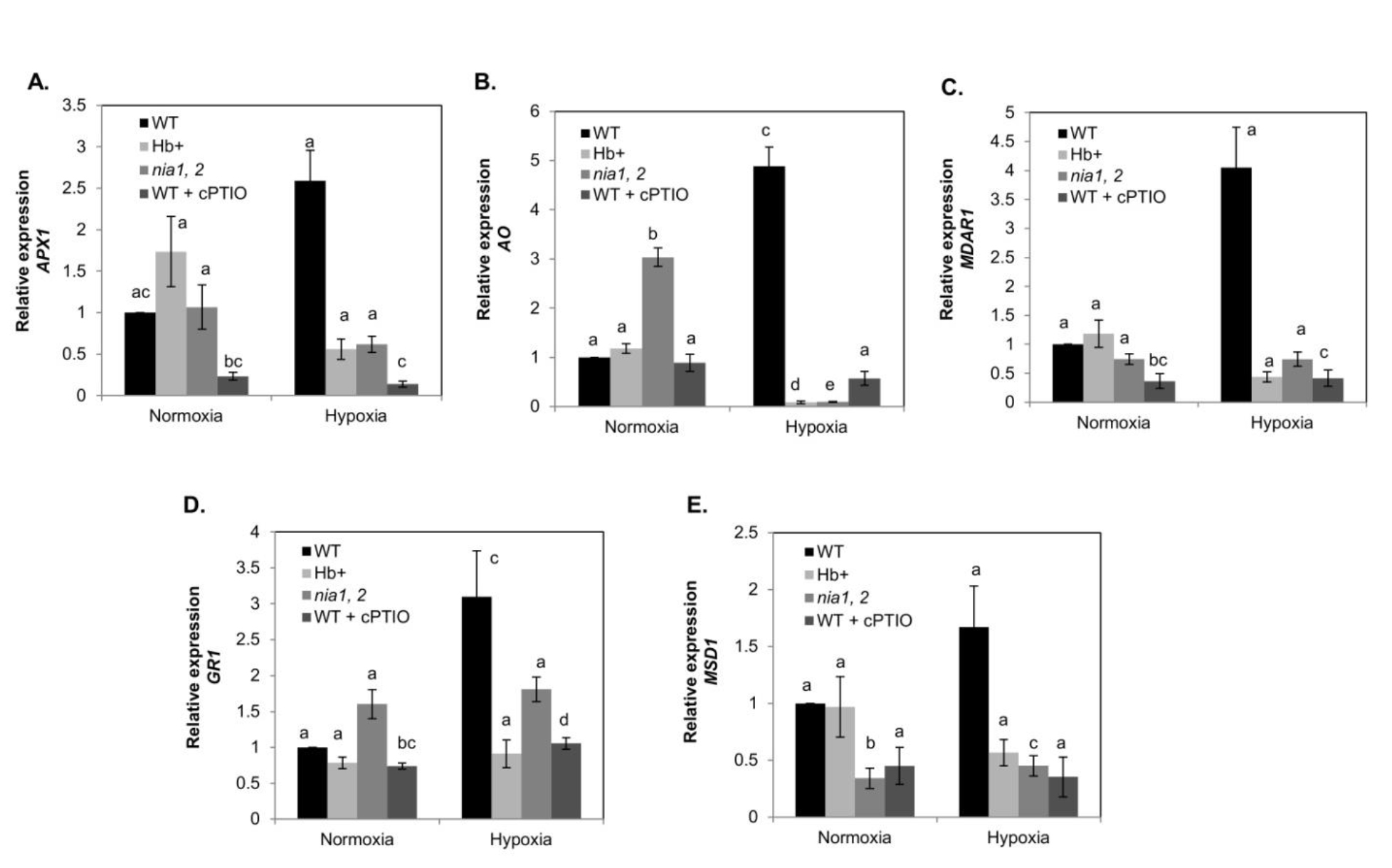
Relative expression of genes involved in antioxidant pathways: (A) *APX1* (*ASCORBATE PEROXIDASE-1*; AT1G07890), (B) *ASCORBATE OXIDASE* (*L-ASCORBATE OXIDASE;* AT5G21100), (C) *MDAR1* (*MONODEHYDROASCORBATE REDUCTASE-1*; AT3G52880), (D) *GR1 (GLUTATHIONE-DISULFIDE REDUCTASE*; AT3G24170) and (E) *MSD1 (MANGANESE SUPEROXIDE DISMUTASE 1*; AT3G10920) in wild-type (WT-black bars), non-symbiotic hemoglobin-1 over-expression lines (Hb+- light grey bars), *nia* double mutants (*nia1,2*– grey bars) and cPTIO pre-treated WT plants (dark grey bars) under normoxia and hypoxia-24h (0.8% O2). Values are means (n=3) ± SE. Statistical significance between normoxia and hypoxia-24h treatments is evaluated by the analysis of variance (ANOVA) followed by different letters indicating significant difference according to Tukey HSD at p<0.01.

**Fig 7:**
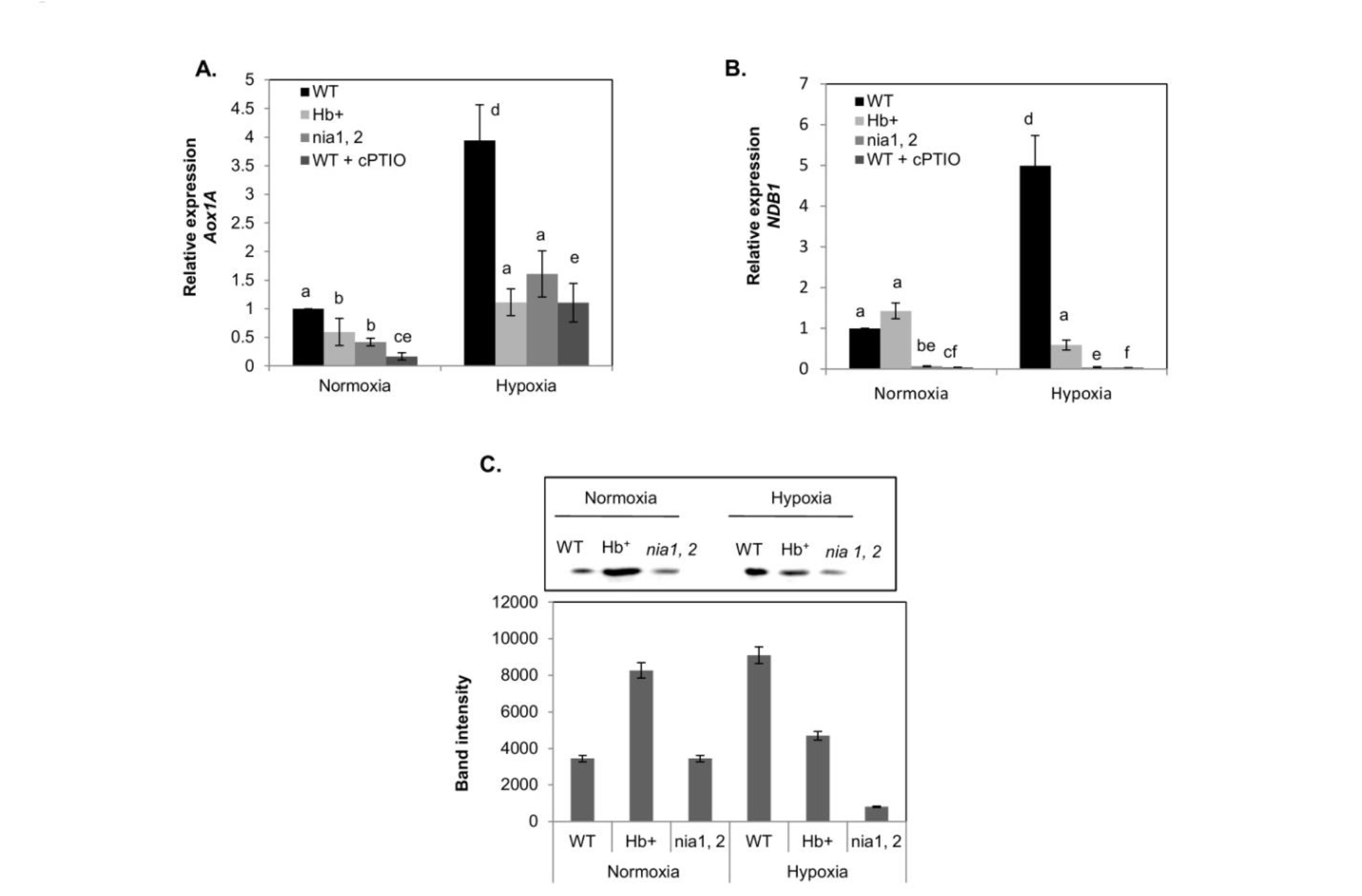
Relative expression of genes in mitochondrial electron transport chain: (A) *AOX1A* (*ALTERNATIVE OXIDASE 1A*; AT3G22370) and (B) *NDB1 (NAD(P)H DEHYDROGENASE B1*; AT4G28220) in wild-type (WT– black bars), non-symbiotic hemoglobin-1 over-expression lines (Hb+- light grey bars), *nia* double mutants (*nia1,2*-grey bars) and cPTIO pre-treated WT plants (dark grey bars) under normoxia and hypoxia-24h (0.8% O2). Values are means (n=3) ± SE. A significant difference between all treatments is analyzed by t-test at P<0.001 (***), P<0.01 (**) and P<0.05 (*) with WT-normoxia as a control. (C) Detection of AOX1A protein abundance in Arabidopsis seedlings (WT, Hb^+^ and *nia1,2* mutant) subjected to hypoxia. Upper panel showing protein bands and lower panel showing band intensity measured by Image J. Blot is representative of two independent experiments.

**Fig 8:**
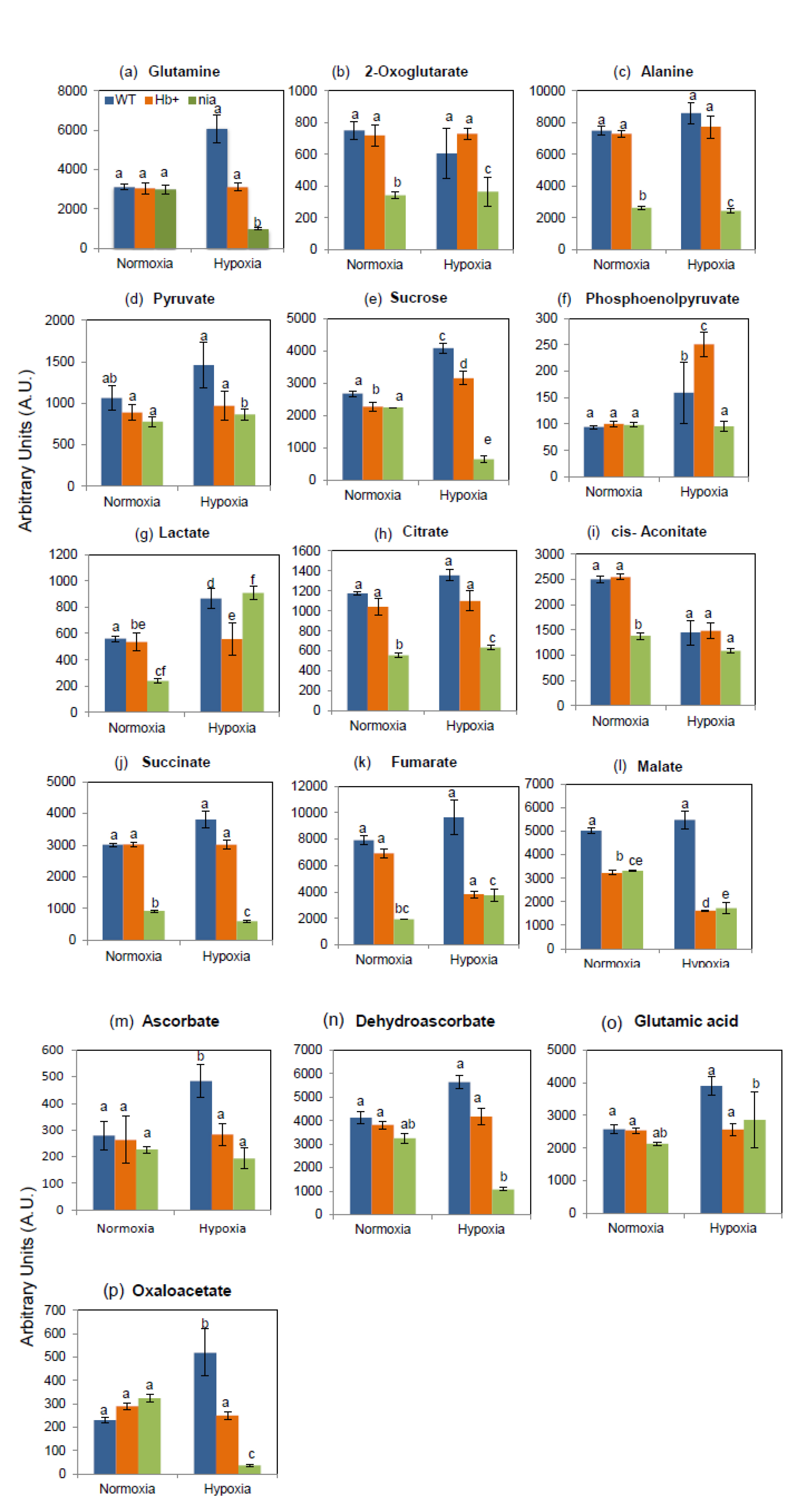
Data for metabolic intermediates of antioxidants, carbon and nitrogen metabolism in the total seedlings of WT (blue bars), Hb+ (orange bars), *nia1,2* (green bars) mutants under normoxia and hypoxia-24h. Quantities are in arbitrary units specific to internal standards for each quantified metabolite and normalized to n=5 for each genotype. Bars represent the SE (n=5). Statistical significance between normoxia and hypoxia-24h treatments is evaluated by the analysis of variance (ANOVA) followed by different letters indicating significant difference according to Tukey HSD at p<0.01.

## Discussion

Oxygen crisis can occur to the plants by many ways during their life cycle either via extreme environmental effects such as flooding and waterlogging or they also face low oxygen stress during their development (Geigenberger, 2003). Several metabolic, anatomical and physiological alterations take place during such stress (Geigenberger *et al*., 2000; Bailey-Serres *et al*., 2012). Here, we studied how the genes involved in these processes are influenced by NO to unlock the potential role of hypoxia-induced NO production.

In this study, we used both transgenic approach and pharmacological approach to see the effect of decreased levels of NO on gene induction. We found that most of the changes observed in Hb+ and *nia* mutant under hypoxia also matched with cPTIO-grown plants suggesting that the observed changes in gene expression under hypoxia are mostly due to reduced levels of NO.

Previously, it was reported that NO production increases during hypoxia/anoxia in various plant species (Gupta *et al*., 2005; Planchet *et al*., 2005) but no attempts were made so far to check the role of the hypoxia-induced NO in gene expression. As expected, NO production was reduced in Hb+ and *nia1,2* mutant as shown by DAF-FM fluorescence (Fig 1A&B). This fluorescence was eliminated by cPTIO suggesting that it is attributed to NO (Fig 1A&B). NR is the major enzyme that is involved in production of NO under hypoxia either directly in the side reaction or by supplying nitrite which is further reduced to NO by mitochondria, therefore, we measured *NIA1* expression. The observed increased expression of *NIA1* in WT under hypoxia suggests that the induction of this gene could contribute to NO synthesis in WT. In Hb+ line and cPTIO-grown WT, there is less pronounced induction of *NIA1* suggesting that scavenging of NO affects the expression of *NIA1,* which further suggests that NO generated under hypoxia amplifies *NIA1* expression. This observation was strengthened by the increased NR activity in WT under hypoxia (Fig 2F).

Expression of the gene *GLN1* encoding glutamine synthetase and the level of glutamine are increased under hypoxia in WT suggesting that the induction of *GLN1* by NO can affect glutamine production positively (Fig 2B & Fig 8a). *2- OXOGLUTARATE DEHYDROGENASE* (*OGDH*) gene encodes the enzyme that links carbon and nitrogen metabolism and thus determines the metabolic flux through the TCA cycle (Tretter & Adam-Vizi, 2007).The decreased level of expression of *OGDH* under hypoxia in WT, Hb+ and *nia1,2* mutant suggests that hypoxic stress supresses *OGDH*. Though this gene is downregulated under hypoxia, its expression was slightly higher in WT suggesting that NO can regulate this gene under hypoxia to maintain metabolic flux of the TCA cycle. Induction of *OGDH* under hypoxia in WT may represent an indirect effect mediated by NO. The limited oxidation of NAD(P)H under hypoxia in mitochondria or inhibition of aconitase by NO both lead to the increased citrate accumulation (Gupta *et al.,* 2012). The accumulated citrate can directly induce *OGDH* to convert 2-oxoglutarate (α- ketoglutarate) to succinyl-CoA and CO2 during the TCA cycle operation under hypoxia to increase metabolic flux.

Another gene encoding the amino acid metabolizing enzyme ferredoxin-dependent glutamate synthase (Fd-GOGAT) *GLU2* was upregulated suggesting that nitrate assimilation pathways genes are induced in the presence of NO. Glutamate synthase utilizes 2-OG to produce glutamate. Our metabolic analysis revealed the reduced levels of 2-OG in WT whereas slight increase of 2-OG in Hb+ under hypoxia suggests active utilization of 2-OG in low oxygen conditions (Fig 8b). Previously, it was shown that alanine aminotransferase plays a role in hypoxia tolerance (Rocha *et al*., 2010a,b; Miyashita *et al*., 2007) via the direction of glycolytic flux towards alanine synthesis with concomitant operation of the incomplete TCA cycle. Alanine aminotransferase plays a role in GABA pathway which is known to regulate pH under hypoxia (Reid *et al.,* 1985). Our results demonstrate that under hypoxia, *AlaAT1* and alanine levels (Fig 2C & 8c) are increased in WT, whereas no induction was observed in *nia1,2* mutant.

Under normoxic conditions, most of the ATP is generated via oxidative phosphorylation but under hypoxic condition, glycolytic ATP is very crucial due to the impairment of the function of electron transport chain to generate ATP (Gupta *et al.,*2009). Previously, it was shown that under anoxia nitrite reduction to NO aids the production of ATP (Stoimenova *et al*., 2007) but, it is not known how the genes of fermentative pathway are modulated by NO. Sucrose synthase provides carbon skeletons for glycolysis. The energetically favourable degradation of sucrose via sucrose synthase has shown to have advantage during hypoxic stress especially when invertase expression is down-regulated and sucrose synthase expression is enhanced (Bologa *et al*., 2003; Narsai *et al*., 2011). Here, our results show that similar trend where *SUCROSE SYNTHASE* (*SUS1*) transcript levels have upregulated 4-fold under hypoxia in WT whereas in Hb+ and *nia1,2* mutant the increase was not prominent under hypoxia. Out of all glycolytic enzymes, pyruvate kinase plays an important role in regulation of glycolysis. This enzyme catalyzes the terminal reaction of the glycolytic pathway by converting ADP and PEP to ATP and pyruvate (Podesta *et al*., 1992), our results show that *PYRUVATE KINASE* (*PK*) has elevated expression in WT under hypoxia and in Hb+ and *nia1,2* mutant its expression was not induced under hypoxia showing the potential requirement of NO to induce this gene.*PK* induction also correlated with pyruvate levels (Fig 8d).

*LACTATE DEHYDROGENASE* (*LDH*) that plays an important role in fermentation has higher expression under hypoxia in WT, again pressing the importance of NO in fermentation. Lactate levels also increased under hypoxia in WT and reduced levels were found in Hb+ and *nia* mutant pointing the importance of NO in regulation of fermentation (Fig 8g). *PYRUVATE DECARBOXYLASE* (*PDC*) converts pyruvate to acetaldehyde which is crucial for the production of ethanol via *alcohol dehydrogenase* (ADH) and for regeneration of NAD^+^ in the later step. Expression analysis revealed that these two genes are 2.5- and 5-fold upregulated in WT under hypoxia suggesting that NO is indeed a major player in regulating fermentative genes under hypoxia. A strong increase of ADH activity in WT and mild increase in Hb+ and *nia* mutant (Fig 3F) suggests that NO modulates not only gene expression but also may modulate the activity of ADH. These results clearly show that NO plays a crucial role in regeneration of NAD^+^ under hypoxia by upregulating these genes.

The increased expression of *ADH1*, *LDH* and *PDC* in WT as compared to Hb+*, nia1,2* under hypoxia (Fig 3) is probably mediated by the induction of group VII ERF transcription factors (Hinz *et al.,* 2010; Licausi et al., 2011). Since NO and ROS both are differentially produced in WT, Hb+, *nia1,2* mutant (Fig 1) the interplay and ratios of NO and ROS might play role in physiological regulation of these genes involved in fermentation under hypoxia. Under hypoxia, RAP2.12 is released from plasma membrane and accumulates in nucleus (Licausi *et al.,* 2011). ROS is mainly generated by mitochondria and plasma membrane NADPH oxidase. The ROS generation by these enzymes also depends on oxygen concentration and NO (Vergara *et al.,* 2012, Wany *et al.,* 2017). Hence, NO production, oxygen and ROS are interlinked and the outcome of these interplay can determine the induction of group VII ERFs. *RAP2.12 and RAP2.2* is also known activate *ADH1* (Papdi *et al.,* 2008; Hinz *et al.,* 2010). The slight induction of *ADH1* in WT than Hb+ and *nia1,2* mutants under hypoxia is probably due to increased production of NO via nitrite dependent pathway but slightly lesser induction of this gene in Hb+ and *nia1,2* mutant than WT correlated with *RAP2.2 expression* (Fig 4C). Although there is increased expression of *RAP2.2* under hypoxia, the slight difference in induction may be due to a differential production of NO and ROS (Fig 1B&C). In WT an increased accumulation of NO is observed, whereas in Hb+ and *nia1,2* mutants the reduced production of NO is seen. However, the increased production of ROS can be seen in Hb+ and *nia1,2* mutants as compared to WT suggesting that, apart from NO, the generation of ROS may play role in induction of *RAP2.2 and ADH1.* A similar trend was observed for *RAP2.12* suggesting that the same mechanism applies to the induction of *RAP2.2* and *ADH1* (Fig 4D). So far no attempts were made to find out the exact ratio of NO and ROS to determine the fate of group VII ERFs. Differential induction of these genes in differential background of ROS and NO observed in WT, Hb+ and *nia1,2* suggests that there is a link between NO and ROS mediated by phytohormones under hypoxia which can explain the induction of group VII ERFs. Experiments using ROS scavenging enzymes in less NO background can unlock the mechanism of interplay between ROS and NO in determining stability of ERFs.

Our results also revealed that the *ATE1, ATE2* gene was down-regulated under hypoxia, and the expression of these genes were suppressed in *ate 1-2* and *prt 6-5* mutants. However, further experiments will be needed in order to define the exact mechanisms by which these factors interact in the presence or absence of NO. We found that WT survived better than Hb^+^, *nia12* mutant suggesting that presence of NO is important for survival of plants under hypoxia (data not shown). However we cant exclude the interaction beteen the NO-producing and scavenging reactions with phytohormones which might collectively contribute for survival under hypoxia.

Analysis of the genes encoding TCA cycle provides some new insights into the additional functions of NO. Oxaloacetate levels are elevated under hypoxia in WT (Fig 8p). *CITRATE SYNTHASE* (*CS4*) gene was highly expressed (nearly 4-fold) under hypoxia in WT. Previously, it was shown that citrate is essential for the induction of *AOX1A* under hypoxia (Gupta *et al*., 2012). Our results further revealed that *CS4* could be partly responsible for the increased citrate levels under hypoxia to induce AOX (Fig 7A & Fig 8h). The expression of *ACONITASE (ACO2)* increased under hypoxia and reduced in Hb+ and *nia 1,2* mutant line. Previously, it was shown that NO inhibits the aconitase enzyme (Navarre *et al.,* 2000). This process was shown to have a protective effect against the additional oxidative stress by acting as a reversible ‘circuit breaker’ (Gardner *et al*., 1991; Gardner *et al*., 1997). Our results indicate that NO can induce the expression of aconitase gene under hypoxia, while the inhibition of aconitase enzyme is attributed to its direct biochemical effect (Gupta *et al.,* 2012). The *nia1,2* mutant exhibited the reduced levels of AOX despite of producing higher ROS than WT suggesting that NO has stronger effect on AOX induction than ROS (Fig 7C). *SUCCINATE DEHYDROGENASE* (*SDH2*) and *FUMARASE* (*FUM1*) gene expression also increased under hypoxia in WT suggesting that NO could contribute to the increased TCA cycle flux via upregulation of succinate dehydrogenase and fumarase genes. Most of the metabolites in fermentative metabolism and TCA cycle have increased under hypoxia suggesting correlation between gene expression and metabolite levels (Fig 8).

Examination of expression of the genes involved in the ascorbate-glutathione cycle and ROS metabolism has revealed many new insights about the importance of hypoxia-induced NO. Expression analysis of *GLUTATHIONE REDUCTASE* (*GR1*), *ASCORBATE PEROXIDASE* (*APX1*) and *MONODEHYDROASCORBATE REDUCTASE-1* (*MDAR1)* revealed that NO can modulate most of the ascorbate-glutathione cycle genes to protect from harmful levels of ROS (Fig 6). Previously, it was shown that ascorbate acts as a primary compound in the reduction of metHb under hypoxic and anoxic conditions (Igamberdiev *et al.,* 2006). The induction of genes involved in the ascorbate-glutathione cycle by NO may help in accelerating Hb-NO cycle to increase hypoxic ATP production. High concentrations of ascorbate in plant cells facilitates scavenging of superoxide (O2-) moreover, ascorbate peroxidase is also involved in H2O2 scavenging and reduction of NO2- to NO (Alegria *et al.,* 2004). The ability of plants under hypoxia to synthesize ascorbate (Fig 8m) and induction of genes involved in the ascorbate-glutathione cycle (Fig 6) may be one of the reasons for production of reduced peroxynitrite under hypoxia (Vishwakarma *et al*., 2018) despite of higher NO production by the mitochondrial COX and AOX (Gupta *et al.,* 2005; Vishwakarma *et al.,* 2018).

*MANGANESE SUPEROXIDE DISMUTASE* (*MSD1*) gene was upregulated under hypoxia in WT suggesting that the produced NO could induce MSD. Therefore, the observed increase in *MSD1* expression can be explained by the increased NO and slightly decreased ROS levels under hypoxia in WT (Fig 1C). ROS act as signals under hypoxia (Baxter-Burrel *et al.*, 2002), but a tight regulation of ROS is important for inducing signalling cascade (Mittler *et al.,* 2011). AOX plays an important role in stress tolerance by accepting excess of electrons from the mitochondrial ubiquinol pool to reduce ROS. We found that under hypoxia, WT plants exhibit the increased expression of *AOX1A* suggesting that NO rather than ROS can influence *AOX1A* gene expression under hypoxia. But under normoxia, despite of the decreased levels of *AOX1A* transcripts in Hb+, the protein levels were very high. This increased levels of the AOX protein in Hb+ are probably due to higher levels of ROS (Fig 1C). Previously, it was shown that under hypoxia, the electron channeling via the external mitochondrial NAD(P)H dehydrogenases (NDB-type) can reduce nitrite to NO (Stoimenova *et al*., 2007). In our experiments, we found that *NDB1* expression exhibited the increase under hypoxia whereas in Hb+ its expression is downregulated suggesting that NO can also induce *NDB1* for oxidizing extra NAD(P)H formed under hypoxia.

## Conclusions

In this study, we investigated the impact of hypoxia on induction of the genes involved in fermentation, TCA cycle, nitrogen metabolism and ascorbate-glutathione cycle (Figure S1). Our study revealed that imposing hypoxia leads to alterations in the levels of NO and ROS. These alterations are either directly mediated by NO or NO-dependent regulation of abscisic acid, ethylene and other hormones biosynthesis, which in turn may cause the observed alterations in gene expression. The increase of NO and ROS under hypoxia was observed in the wild type plants while the reduced levels of NO and increased levels of ROS were observed in Hb+ and *nia1,2* mutant plants. Strikingly, it was found that the fermentative pathway enzymes such as alcohol dehydrogenase (ADH) and pyruvate decarboxylase (PDC) are modulated by hypoxia-induced NO and ROS. We also found that NO can induce various TCA cycle genes and metabolites in order to accelerate its capacity to regenerate reducing equivalents under hypoxia. ROS detoxifying genes involved in the ascorbate-glutathione cycle and other ROS-scavenging pathways are highly induced by NO under hypoxia. Overall our study revealed that hypoxia-induced NO plays an important role in modulation of genes and metabolites involved in carbon, nitrogen and antioxidant metabolism which are crucial for survival of plants under low oxygen stress.

## Author contributions

KJG designed experiments, AW, YB, SP, APV, AK, PS, PKP, AKG conducted experiments, analysed data and prepared graphs. KJG wrote the manuscript.

## Acknowledgements

KJG was supported by DBT Ramalingaswami Fellowship and IYBA grant award from Department of Biotechnology, Govt of India. AW, SP was supported by National Postdoctoral Fellowship from SERB, India. PKP and PS was supported by UGC JRF and SRF fellowships. We thank Kim Hebelstrup for providing non-symbiotic haemoglobin overexpressing line. We acknowledge the central instrumental facility of NIPGR for technical support.

## References

Abiko T, Kotula L, Shiono K, Malik AI, Colmer TD, Nakazono M. 2012. Enhanced formation of aerenchyma and induction of a barrier to radial oxygen loss in adventitious roots of *Zea nicaraguensis* contribute to its waterlogging tolerance as compared with maize (*Zea mays* ssp. mays). Plant, Cell and Environment 35, 1618–1630.

Alegria AE, Sanchez S, Quintana I. 2004. Quinone-enhanced ascorbate reduction of nitric oxide: role of quinone redox potential. Free Radical Research 38, 1107–1112.

Bailey-Serres J, Lee SC, Brinton E. 2012. Waterproofing Crops: Effective Flooding Survival Strategies. Plant Physiology 160, 1698–1709.

Baxter-Burrell A, Yang Z, Springer PS & Bailey-Serres J. 2002. RopGAP4-dependent Rop GTPase rheostat control of Arabidopsis oxygen deprivation tolerance. Science 296, 2026–2028.

Benamar A, Rolletschek H, Borisjuk L, Avelange-Macherel MH, Curien G, Mostefai HA, Andriantsitohaina R, Macherel D. 2008. Nitrite-nitric oxide control of mitochondrial respiration at the frontier of anoxia. Biochimica et Biophysica Acta 1777, 1268–1275.

Bologa KL, Fernie AR, Leisse A, Loureiro ME, Geigenberger P. 2003. A bypass of sucrose synthase leads to low internal oxygen and impaired metabolic performance in growing potato tubers. Plant Physiology 132, 2058–2072.

Borisjuk L, Macherel D, Benamar A, Wobus U, Rolletschek H. 2007. Low oxygen sensing and balancing in plant seeds – a role for nitric oxide. New Phytologist 176, 813–823.

Bradford, M. M. (1976). A rapid and sensitive method for the quantitation of microgram quantities of protein utilizing the principle of protein dye binding. Analytical Biochemistry, 72, 248–254.

Bright J, Desikan R, Hancock JT, Weir IS, Neill SJ. 2006. ABA-induced NO generation and stomatal closure in Arabidopsis are dependent on H2O2 synthesis. Plant J 45, 113–122.

Colmer TD, Voesenek LACJ. 2009. Flooding tolerance: suites of plant traits in variable environments Functional Plant Biology 36, 665–681.

Corpas FJ, Hayashi M, Mano S, Nishimura M, Barroso JB. 2009. Peroxisomes are required for in vivo nitric oxide accumulation in the cytosol following salinity stress of Arabidopsis plants. Plant Physiology 151, 2083–2094.

Cuadros-Inostroza A, Caldana C, Redestig H, Kusano M, Lisec J, Peña-Cortés H, Willmitzer L, Hannah MA. 2009. Target Search – a Bioconductor package for the efficient preprocessing of GC-MS metabolite profiling data, BMC Bioinformatics, 10 428.

Drew MC. 1997. Oxygen deficiency and root metabolism: injury and acclimation under hypoxia and anoxia. Annual Review of Plant Physiology and Plant Molecular Biology 48, 223–250.

Fox TC, Kennedy RA, Rumpho ME. 1994. Energetics of plant growth under anoxia: metabolic adaptations of *Oryza sativa* and *Echinochloa phyllopogon*. Annals of Botany 74, 445–455.

Gardner PR, Costantino G, Szabo C, Salzman AL. 1997. Nitric oxide sensitivity of the aconitases. Journal of Biological Chemistry 272, 25071–25076

Geigenberger P, Fernie AR, Gibon Y, Christ M, Stitt M. 2000. Metabolic activity decreases as an adaptive response to low internal oxygen in growing potato tubers. Journal of Biological Chemistry 381, 723–740.

Geigenberger P. 2003. Response of plant metabolism to too little oxygen. Current Opinion in Plant Biology 6, 247–256.

Gibbs DJ, Lee SC, Isa NM, Gramuglia S, Fukao T, Bassel GW, Correia CS, Corbineau F, Theodoulou FL, Bailey-Serres J and Holdsworth MJ. 2011. Homeostatic response to hypoxia is regulated by the N-end rule pathway in plants. Nature 479, 415–418.

Gupta KJ, Fernie AR, Kaiser WM, van Dongen JT. 2011) On the origins of nitric oxide. Trends in Plant Science 16:160–168.

Gupta KJ, Hebelstrup KH, Kruger NJ, Ratcliffe RG. 2014. Nitric oxide is required for homeostasis of oxygen and reactive oxygen species in barley roots under aerobic conditions. Molecular Plant 7, 747–750.

Gupta KJ, Shah JK, Brotman Y, Jahnke K, Willmitzer L, Kaiser WM, Bauwe H, Igamberdiev AU. 2012. Inhibition of aconitase by nitric oxide leads to induction of the alternative oxidase and to a shift of metabolism towards biosynthesis of amino acids. Journal of Experimental Botany, 63, 1773–1784.

Gupta KJ, Stoimenova M, Kaiser WM. 2005. In higher plants, only root mitochondria, but not leaf mitochondria reduce nitrite to NO, in vitro and in situ. Journal of Experimental Botany 56, 2601–2609.

Gupta KJ, Zabalza A, van Dongen JT. 2009. Regulation of respiration when the oxygen availability changes. Physiologia Plantarum 137, 383–391.

Gardner PR, Fridovich I. 1991. Superoxide sensitivity of the *Escherichia coli* aconitase. Journal of Biological Chemistry 266, 19328–19333.

Hattori Y, Nagai K, Furukawa S, et al. 2009. The ethylene response factors SNORKEL1 and SNORKEL2 allow rice to adapt to deep water. Nature 460, 1026–1030.

Hinz M, Wilson IW, Yang J, Buerstenbinder K, Llewellyn D, Dennis ES, Sauter M, and Dolferus R. 2010. Arabidopsis RAP2.2: an ethylene response transcription factor that is important for hypoxia survival. Plant Physiology 153, 757–772.

Hebelstrup KH, Jensen EO. 2008. Expression of NO scavenging hemoglobin is involved in the timing of bolting in *Arabidopsis thaliana*. Planta 4, 917–27.

Huang B, Johnson JW, NeSmith S, Bridge DC. 1994. Growth, physiological and anatomical responses of two wheat genotypes to waterlogging and nutrient supply. Journal of Experimental Botany 45, 193–202.

Igamberdiev AU, Bykova NV, Hill RD. 2006. Scavenging of nitric oxide by barley hemoglobin is facilitated by a monodehydroascorbate reductase-mediated ascorbate reduction of methemoglobin. Planta 223, 1033–1040.

Igamberdiev AU, Hill RD. 2004. Nitrate, NO and haemoglobin in plant adaptation to hypoxia: an alternative to classic fermentation pathways. Journal of Experimental Botany 55, 2473–2482.

Kopka J, Schauer N, Krueger S, et al. 2005: the Golm Metabolome Database, Bioinformatics.15, 1635–1638.

Lisec J, Schauer N, Kopka J, Willmitzer L, Fernie AR. 2006. Gas chromatography mass spectrometry-based metabolite profiling in plants, Nature Protocols 1, 387–396.

Licausi F, Kosmacz M, Weits DA, Giuntoli B, Giorgi FM, Voesenek LA, Perata P, and van Dongen JT. 2011. Oxygen sensing in plants is mediated by an N-end rule pathway for protein destabilization. Nature 479, 419–422.

Mergemann H, Sauter M. 2000. Ethylene induces epidermal cell death at the site of adventitious root emergence in rice. Plant Physiology 124, 609–614.

Mittler R, Vanderauwera S, Suzuki N, Miller G, Tognetti VB, Vandepoele K, Gollery M, Shulaev V, Van Breusegem F. 2011. ROS signaling: the new wave? Trends in Plant Science 16, 300–309.

Miyashita Y, Dolferus R, Ismond KP, Good AG. 2007. Alanine aminotransferase catalyses the breakdown of alanine after hypoxia in *Arabidopsis thaliana*. Plant Journal 49, 1108–1121.

Mur LA, Mandon J, Persijn S, Cristescu SM, Moshkov IE, Novikova GV, Hall MA, Harren FJ, Hebelstrup KH, Gupta KJ. 2013. Nitric oxide in plants: an assessment of the current state of knowledge. AoB Plants. 5, pls052.

Mustroph A, Lee SC, Oosumi T, Zanetti ME, Yang H, Ma K, Yaghoubi-Masihi A, Fukao T, Bailey-Serres J. 2010. Cross-kingdom comparison of transcriptomic adjustments to low-oxygen stress highlights conserved and plant-specific responses. Plant Physiology 152, 1484–1500.

Navarre DA, Wendehenne D, Durner J, Noad R, Klessig DF. 2000. Nitric oxide modulates the activity of tobacco aconitase. Plant Physiology 122, 573–582.

Papdi C, Abraham E, Joseph MP, et al. 2008.Functional identification of Arabidopsis stress regulatory genes using the controlled cDNA overexpression system. Plant Physiology 147, 528–542.

Planchet E, Gupta KJ, Sonoda M, Kaiser WM. 2005. Nitric oxide emission from tobacco leaves and cell suspensions: rate-limiting factors and evidence for the involvement of mitochondrial electron transport. Plant Journal 41, 732–743.

Podesta FE, Plaxton WC. 1992. Plant cytosolic pyruvate kinase: a kinetic study. Biochimica et Biophysica Acta 1160, 213–220.

Ricard B, Couee I, Raymond P, Saglio PH, Saint-Ges V, Pradet A. 1994. Plant metabolism under hypoxia and anoxia. Plant Physiology and Biochemistry 32, 1–10.

Rocha M, Sodek L, Licausi F, Hameed MW, Dornelas MC, van Dongen JT. 2010a. Analysis of alanine aminotransferase in various organs of soybean (*Glycine max*) and in dependence of different nitrogen fertilizers during hypoxic stress. Amino Acids 39, 1043–1053.

Reid RJ, Loughman BC, Ratcliffe RG. 1985. ^31^P NMR measurements of cytoplasmic pH changes in maize root tips. Journal of Experimental Botany 36, 889–897.

Rocha M, Licausi F, Araujo WL, Nunes-Nesi A, Sodek L, Fernie AR, van Dongen JT. 2010b. Glycolysis and the tricarboxylic acid cycle are linked by alanine aminotransferase during hypoxia induced by waterlogging of *Lotus japonicus*. Plant Physiology 152, 1501–1513.

Rolletschek H, Borisjuk L, Koschorreck M, Wobus U, Weber H. 2002. Legume embryos develop in a hypoxic environment. Journal of Experimental Botany 53, 1099–1107.

Royo B, Moran JF, Ratcliffe RG, Gupta KJ. 2015. Nitric oxide induces the alternative oxidase pathway in Arabidopsis seedlings deprived of inorganic phosphate. Journal of Experimental Botany 66, 6273–2680.

Rumer S, Gupta KJ, Kaiser WM. 2009. Oxidation of hydroxylamines to NO by plant cells. Plant Signaling & Behavior 4, 853–855.

Sakihama Y, Nakamura S, Yamazaki H.2002. Nitric oxide production mediated by nitrate reductase in the green alga *Chlamydomonas reinhardtii*: an alternative NO production pathway in photosynthetic organisms. Plant and Cell Physiology 43, 290–297.

Stöhr C, Strube F, Marx G, Ullrich WR, Rockel P.2001. A plasma membrane-bound enzyme of tobacco roots catalyses the formation of nitric oxide from nitrite. Planta 212, 835–841.

Stoimenova M, Igamberdiev AU, Gupta KJ, Hill RD. 2007. Nitrite driven anaerobic ATP synthesis in barley and rice root mitochondria. Planta 226, 465–474.

Steinfath M, Groth D, Lisec J, Selbig J. 2008. Metabolite profile analysis: from raw data to regression and classification, Physiologia Plantarum 132, 150–161.

Tretter L, Adam-Vizi V. 2007. Uncoupling is without an effect on the production of reactive oxygen species by in situ synaptic mitochondria. Journal of Neurochemistry 103, 1864–1871.

Tun NN, Santa-Catarina C, Begum T, Silveira V, Handro W, Floh EI, Scherer GF. 2006. Polyamines induce rapid biosynthesis of nitric oxide (NO) in Arabidopsis thaliana seedlings. Plant and Cell Physiology 47, 346–354.

Toufighi, K, Brady SM, Austin R, Ly E, Provart, NJ. 2005. The Botany Array Resource: e-Northerns, Expression Angling, and promoter analyses. Plant Journal 43, 153–163.

Vergara R, Parada F, Rubio S, Perez FJ. 2012. Hypoxia induces H2O2 production and activates antioxidant defence system in grapevine buds through mediation of H_2_O_2_ and ethylene. Journal of Experimental Botany 63, 4123–4131.

Vishwakarma A, Kumari A, Mur LAJ, Gupta KJ. 2018. A discrete role for alternative oxidase under hypoxia to increase nitric oxide and drive energy production. Free Radical Biology and Medicine 122, 40–51.

Wany A, Kumari A, Gupta KJ. 2017 Nitric oxide is essential for the development of aerenchyma in wheat roots under hypoxic stress. Plant Cell and Environment 40, 3002–17.

Winkel A, Colmer TD, Pedersen O. 2011. Leaf gas films of *Spartinaanglica* enhance rhizome and root oxygen during tidal submergence. Plant Cell and Environment 34, 2083–2092.

Zabalza A, Van Dongen JT, Froehlich A, et al. 2009. Regulation of respiration and fermentation to control the plant internal oxygen concentration. Plant Physiology 149, 1087–1098.

